# mastR: Marker Automated Screening Tool for multi-omics data

**DOI:** 10.1101/2024.04.24.590691

**Authors:** Jinjin Chen, Ahmed Mohamed, Dharmesh D. Bhuva, Melissa J. Davis, Chin Wee Tan

## Abstract

**Motivation:** Biomarker discovery and utilization is important and offers insight into potential underlying mechanisms of disease. Existing marker identification methods primarily focus on single cell RNA sequencing (scRNA-seq) data, with no specific automated methods designed to learn from the bulk RNA-seq data. Furthermore, when adapting scRNA-seq methods to bulk RNA-seq, the background expressions of non-targeted cell types are not accounted for. Here we bridge this gap with an automated marker identification method that works for bulk RNA sequencing data.

**Results:** We developed *mastR*, a novel computational tool for accurate marker identification from omics data. It leverages robust pipelines from *edgeR* and *limma* R/Bioconductor packages, performing pairwise comparisons between groups, and aggregating the results through rank-product-based permutation test. A signal-to-noise ratio approach is implemented to minimize background signals. We assess the performance of a *mastR*-derived NK cell signature against curated published signatures and find our derived signature performs as well if not better than published signatures. We also demonstrate the utility of *mastR* on simulated scRNA sequencing data and provide examples of *mastR* outperforming *Seurat* in marker selection.

**Availability and implementation:** All statistical analyses were carried out using R (version 4.3.0 or higher) and Bioconductor (version 3.17 and higher). *MastR* is available as an R/Bioconductor package with a comprehensive vignette for ease of use (https://bioconductor.org/packages/release/bioc/html/mastR.html) and a guide hosted on GitHub: https://davislaboratory.github.io/mastR/.

## 1. Introduction

Biomarkers are biological features that infer the states of cells, tissues, or individuals, either diseased or healthy. Biomarkers may include molecular features like genes, and proteins which can be used in research and clinical settings to provide insights into disease diagnosis, prognosis and treatment. In recent years, biomarkers have been identified through various ∼omics approaches, including transcriptomics, proteomics, and metabolomics, providing an overview of the molecular landscape of the system being studied (Lawlor, et al., 2009; Rodrigues, et al., 2016; Wang and Yu, 2013). This influx of omics data has advanced the development of computational and bioinformatics methods to identify biological markers, providing the opportunities to accelerate biomarker discovery and thereby facilitating diagnostic and therapeutic developments for various diseases and cancers (Kaur, et al., 2021; Lee and Kim, 2021; Vlachavas, et al., 2021). However, deciphering the background signal of the sample microenvironment from the desired marker signal remains a complex issue to be overcome.

Currently, most marker identification methods focus only on the latest data types such as single-cell RNA sequencing (scRNA-seq) data or spatial data, where the disease signal is easier to separate from the background signal, with no methods looking into using the large amount of bulk RNA sequencing (RNA-seq) data for automated marker identification. To use bulk RNA-seq data for marker identification, there is still a need for either an existing potential marker list (e.g. *MarkerPen* (Qiu, et al., 2021)), or manual selection of differential expression (DE) analysis results from *edgeR*, *limma* (Baran and Dogan, 2023; Dumitrascu, et al., 2021; Fu, et al., 2022; Nelson, et al., 2022) which also involves simple collation of DE genes from different comparisons and datasets. Most of these collations disregard the statistical information from each comparison or dataset analysis, and the statistical information is typically not fully utilised when performing curation. Further, the effects of background RNA expression in the sample microenvironment (tissue or organ) are not considered by these methods, potentially introducing non-specific findings. To ensure accurate, and specific marker identification, the non-disease background expression of a sample must be adjusted for, so the marker-specific signal is not confounded.

To address these considerations, we developed an R/Bioconductor package *mastR* (**M**arkers **A**utomated **S**creening **T**ool in **R**), an automated marker gene identification tool for omics data. It is developed for marker genes selection, employing *edgeR* and *limma* for robust statistical analysis. *MastR* then uses a rank-product-based scoring approach to integrate statistical information of interest across DE comparisons. Finally, *mastR* accounts for background gene expression by explicitly encoding the signal-to-noise ratio (SNR) into the marker selection algorithm. Here we introduce *mastR*, presenting the utility and various applications of the package. We demonstrate that *mastR* is highly accurate, computationally efficient and transferable to several simulated and public datasets, emphasizing its potential research and clinical application in the identification of reliable biological markers.

## 2. Methods

### 2.1 Data

In this study, we accessed publicly available data including all the samples from the DICE (Database of Immune Cell Expression, Expression quantitative trait loci (eQTLs) and Epigenomics) project (https://dice-database.org) (Schmiedel, et al., 2018), colorectal cancer (CRC) RNA-seq data from The Cancer Genome Atlas (TCGA) (Cancer Genome Atlas, 2012), CRC cell line RNA-seq (TPM normalised) data from the Cancer Cell Line Encyclopaedia (CCLE) (https://sites.broadinstitute.org/ccle) (Barretina, et al., 2012), whole transcriptome RNA-seq data of 6 sorted immune cell types of healthy individuals from GSE60424 (depicted as *im_data_6*, (Linsley, et al., 2014)) and a scRNA-seq peripheral blood mononuclear cell (PBMC) data (*pbmc3k.final*) in the Bioconductor R package *SeuratData* (Lab, 2020).

For comparisons of signatures, the published curated natural killer (NK) cell signatures from Crinier et al. (Crinier, et al., 2018) (named as *NK Crinier*), Cursons et al. (Cursons, et al., 2019) (named as *NK Cursons*) and Shembrey et al. (Shembrey, et al., 2022) (named as *NK Shembrey*) were used. All publicly available datasets and signatures are listed in **Table 1**.

**Table 1.** Datasets used in this study.

| Data | Source | Date<br>Version | Reference |
| --- | --- | --- | --- |
| DICE | Database of Immune Cell Expression, Expression quantitative trait loci (eQTLs) and Epigenomics | February 2018 | PMID: 30449622 |
| TCGA-COAD RNA-seq | The UCSC Cancer Genomics Browser | March 2023 | PMID: 22810696 |
| CCLE RNA-seq | Cancer Cell Line Data Repository | May 2023 | PMID: 22460905 |
| GSE60424 | Gene Expression Omnibus | January 2015 | PMID: 25314013 |
| pbmc3k.final | SeuratData | pbmc3k 3.1.4 | https://github.com/satijalab/seurat-data |
| NK Crinier | Crinier et al. | November 2018 | PMID: 30413361 |
| NK Cursons | Cursons et al. | July 2019 | PMID: 31088844 |
| NK Shembrey | Shembrey et al. | January 2023 | PMID: 36685584 |

### 2.2 Data preprocessing

For counts data (i.e. *im_data_6* and simulated data), *mastR* automatically applies the standard *edgeR* (Robinson, et al., 2010) and *limma-voom* (Ritchie, et al., 2015) filtering and normalization pipeline as described in **Methods 2.3.2**. In cases with normalized/processed data (i.e. DICE, CCLE and *pbmc3k.final*), *mastR* conducts gene filtering based on user-defined thresholds (in this study we use *edgeR filterByExpr* function) without normalization, to avoid over correction/normalization. *MastR* also provides conversion for gene identifiers that do not conform to the HGNC (HUGO Gene Nomenclature Committee) symbol nomenclature, to gene symbols (based on the *org.Hs.eg.db* database (Carlson, 2023)) to ensure standardized gene nomenclature.

### 2.3 Methodology for the *mastR* method

Application of the *mastR* workflow involves 4 sections (**Figure 1**): 1) build a markers pool; 2) identify signature of the target group; 3) refine signature by removing the background signal of the sample microenvironment; 4) visualize the resulting signature.

**Figure 1.**
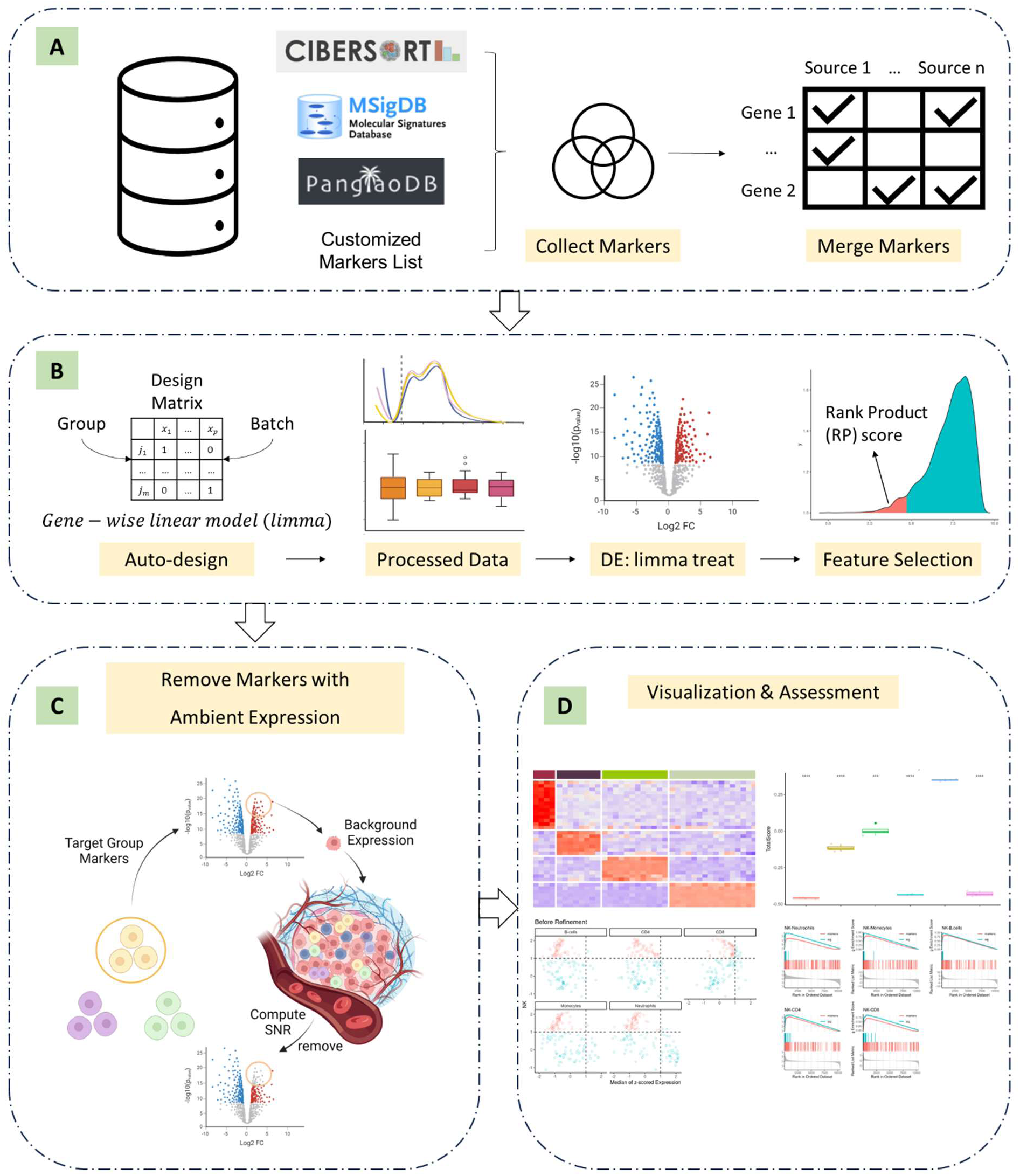
Schematic of the *mastR* workflow. The whole workflow of *mastR* can be divided into 4 main sections: (A) build markers pool; (B) identify signature markers; (C) refine signature by filtering based on background expression and (D) visualize and access signature performance. The *mastR* workflow recommends integrating markers from multiple sources (e.g. PanglaoDB, MSigDB) to form an initial set of markers. *MastR* then generates a design matrix based on the given “Group” and “Batch” factors to be used during data processing and DE analysis. The data processing includes a standard *edgeR* data filtration and normalization pipeline, and a *limma-voom-treat* based linear modeling DE approach conducted to compare the target group with all other groups. *MastR* then computes the marker’s rank product (RP) score based on the ranked product across the DE comparisons and bootstrapped permutation null distribution for further feature selection across multiple comparisons. The selected features will be constrained by the intersection with the initial set of markers. *MastR* allows for filtrating of genes based on the signal-to-noise ratio (SNR) with a background dataset to remove features with inherent expression in a specific context or disease. *MastR* then provides useful visualization functions to assess signature performance.

#### 2.3.1 Build a marker pool

A standard *mastR* pipeline begins by generating a pool of candidate markers. This pool can be compiled either by using the functions in *mastR* or by custom curation and selection of marker genes from other databases or publications. In the former, the R/Bioconductor package *mastR* allows extraction of marker genes specific to immune cell types, relevant pathways, and/or gene sets from existing data sources, which can be retrieved via *get_lm_sig/get_panglao_sig/get_gsc_sig* functions in *mastR*. This includes leukocyte gene signature matrices from CIBERSORT (LM7 (Tosolini, et al., 2017) and LM22 (Newman, et al., 2015)), PanglaoDB (single cell RNA sequencing experiments from mouse and human) (Franzen, et al., 2019), and MSigDB (Molecular Signatures Database) (Liberzon, et al., 2015; Subramanian, et al., 2005) respectively.

LM7 and LM22 are immune cell signature matrices defining 7 and 22 immune cell types, respectively. These matrices are available from CIBERSORT (tool for deconvolving complex tissues based on gene expression profiles) (Newman, et al., 2015). PanglaoDB (Franzen, et al., 2019) is a database of single-cell RNA sequencing experiments that includes marker genes for 25 different immune cell types. MSigDB, or Molecular Signatures Database (Liberzon, et al., 2015), is a collection of annotated gene sets commonly used for pathway analysis.

*MastR* provides *gsc_plot* function to help visualize the overlap of sets of markers. These sets of marker genes can be seamlessly merged as the original pool of markers using the *merge_markers* function in *mastR*, with all marker gene sources saved in the *longDescription* attribute. This merged pool will be used in subsequent analyses. When the markers pool is used in downstream filtration, all the marker genes in the pool will be preserved (as these are determined to be of biological significance) and will not be affected by the filtering step.

#### 2.3.2 Identify signature of the target group

To identify group-specific signatures, *mastR* uses 3 main steps: (a) differential expression analysis (DEA), (b) feature selection to select highly differentially expressed genes (DEGs) based on their rank-product score; and (c) constrain selected genes within the markers pool (**Figure 1B**).

Firstly, DEA is performed using standard *edgeR* (Robinson, et al., 2010) and *limma* (Ritchie, et al., 2015) workflow (i.e., filtering, normalizing, sample weighting and linear modeling). Given the “Group” and “Batch” factors in the data, *mastR* can automatically generate the appropriate design matrix to be used during data filtration, normalization and batch effect correction. Here batch factor is used as fixed effect in linear modelling as it was found the use of batch-corrected data rarely improves the analysis for sparse data, whereas batch covariate modeling improves the analysis for substantial batch effects (Nguyen, et al., 2023). mastR allows for either raw counts or log-normalized data as input, different processing pipelines will be conducted on different types of input. Raw count data is filtered by the *filterByExpr* function in *edgeR*, normalized using the trimmed mean of M-values (TMM) method and analyzed using the *“limma voom with treat”* pipeline. While for log-normalized data, genes are filtered by user-defined thresholds and *“limma trend with treat”* method is employed. In most cases we examined, the standard parameter of log-fold-change (logFC) equal to 0 was used to perform DE analysis, except for the generation of NK cell signature using DICE where a logFC of 1.5 was used.

Secondly, feature selection is conducted to select genes specific to the target group across multiple comparisons. The probability *score*_g_is computed by comparing the rank product score *RP*_g_with permutated random score *rp* from bootstrap approach (Equation (1)-(3)). The common DE genes from *n* − 1 comparisons (where *n* is the total number of groups) are identified and ranked based on the given gene statistics (e.g., p value, adjusted p-value or log fold change) for each comparison. The ranks for each marker gene across all comparisons are log-transformed and summed, before applying a permutation test to bootstrap the null distribution of the random rank product. The resulting marker genes are ordered by *score*_g_ (where smaller values are more significant) and filtered by a selected threshold (by default 0.05).

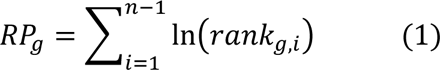

where *rank*_g,i_ is the rank of gene g in *i*^th^ list,

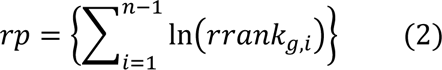

where *rrank*_g,i_ is the shuffled rank of gene g in *i*^th^ list,

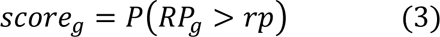

The computation can be summarized as below:

1. Rank the gene statistics in increasing order (decreasing order of ||logFC|| when statistics is logFC) ⇒ *rank*_g,i_: rank of g^th^ gene under i^th^ comparison;
2. Sum log-rank for each gene across comparisons as *RP*_g_: rank product of g^th^ gene;
3. Independently permute statistics value within each comparison relative to gene ID, repeat step (1)–(2) ⇒ 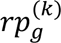: random rank product of g^th^ gene;
4. Repeat step (3) *K* times, form reference null distribution with 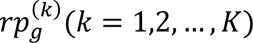;
5. Determine the probability associated with each gene ⇒ *score*_g_. To accommodate analysis with closely similar groups and to improve the specificity for the target group, the threshold filtering based on rank product can be omitted for the target comparison(s) in question by setting parameters “keep.top” and “keep.group”, allowing for more DEGs in the targeted comparison(s).

Thirdly, the identified marker genes are limited to those in the markers pool (i.e. common genes are retained) as the resulting signature. This refinement approach enhances both the discriminative power and the precision of the resulting signature when there is prior knowledge. When the input involves multiple datasets, *mastR* aggregates the individual signature lists identified by each dataset using either a “Robust Rank Aggregation (RRA)”, “union”, or “intersect”.

The aggregation method “RRA” detects marker genes that are consistently ranked higher than stochastically expected under the null hypothesis of uncorrelated inputs and assigns a significance score to each gene (Kolde, et al., 2012). It is suggested for robust gene selection from large number of DEGs. The “union” method is recommended for small size of marker genes identified per dataset; and the “intersect” method, is a simplistic approach, best used in situations characterized by high levels of marker intersection.

*MastR* provides a series of step-by-step functions as well as an integrated wrapper function to implement the above analyses.

#### 2.3.3 Refine signature by accounting for background expression

To avoid background microenvironment confounding effects, *mastR* can further refine the marker genes by filtering out those with ubiquitous expression. *MastR* utilizes an approach which filters out marker genes with low “signal-to-noise ratio” (SNR), which therefore have limited discriminative power between the group of interest and the “background” or “environment” (equations (4-5)). Considering the background expression and signal expression do not always originate from the same batch, and that re-normalizing the entire data is time-consuming and redundant, here to make the sample microenvironment and signal data comparable between batches, the relative expression of the genes within the samples are used to compute SNRs, which makes “signal” and “noise” more comparable across datasets.

Given the widespread assumption that most genes exhibit non-differential expression, the genes expression within each sample should follow a normal distribution denoted as *X*∼*N*(*μ*, *σ*^2^). The 2 parameters of interest, mean (*μ*) and standard deviation (*σ*), can be estimated through maximum likelihood estimation (MLE). For a normal distribution, the likelihood function is the joint probability density function of the data. MLE aims to maximize the logarithm of the likelihood function (equation (4)) to find the best-fitting parameters. Then the percentile (accumulated density) for each gene in each sample can be obtained using the Gaussian cumulative distribution function (CDF) *F*(*x*|μ, σ), and SNR can be computed as outlined in equation (5).

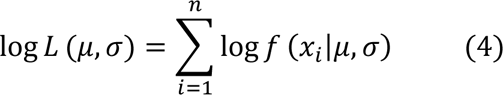

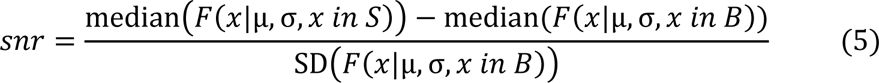

where *x*_i_ are the observed log-transformed gene expression, *n* is the total number of genes, *f* is the probability density function (PDF), *μ* is the mean of the normal distribution, *σ* is the standard deviation of the normal distribution, *snr* is the signal-to-noise ratio of each gene, *S* is the signal dataset and *B* is the background dataset.

This crucial step removes the effect of sample purity for the identified signature markers. By excluding the marker genes with similar expression in the sample microenvironment, the SNR approach ensures only the marker genes with robust and specific expression patterns in the group of interest are retained, leading to a more refined and accurate signature marker list.

#### 2.3.4 Visualize signature performance

*MastR* provides a suite of visualization functions including heatmaps, box plots, scatter plots, rank density plots, and gene set enrichment analysis (GSEA) plots to facilitate the interpretation of the results and enable the evaluation of signature performances across multiple datasets. Details of these visualizations can be found in **Table 2** and described in the vignette (https://davislaboratory.github.io/mastR/articles/mastR_Demo.html).

**Table 2.** Visualization Descriptions.

| Visualization Function | Purpose |
| --- | --- |
| **heatmaps** | Provide an intuitive depiction of the expression levels of individual signature genes across various groups. |
| **box plots** | Assist in evaluating the overall separability of signatures between different groups by representing the distribution of signature scores within each group. |
| **scatter plots** | Allow for the examination of group specificity for each gene and the co-expression patterns between different groups, which can aids in the further refinement of gene selection. |
| **rank density plots** | Visually show the overall expression level of the signature in different groups. |
| **gene set enrichment analysis (GSEA) plots** | Simplify the demonstration of how given signature(s) are enriched in the target group compared to other groups. |

### 2.4 Simulations for assessing performance

#### 2.4.1 Robustness

The robustness of *mastR* method was evaluated using simulated bulk RNA-seq data with varying sample sizes and group proportions. The simulated data was generated as per described by Law *et al*. (Law, et al., 2014) consisting of 100 samples across 4 groups (25 samples per group) with 1000 genes each where 2% are differentially expressed (either up- or down-regulated). The group-specific signatures were identified on the whole simulated dataset using *mastR* first. A subset of the dataset was then generated by random sampling using either a balanced probabilities strategy or an imbalanced probabilities strategy for each group. In each sub-dataset, the new group-specific signatures were identified by *mastR*, and compared to the known simulated DEGs (ground truth) or the signatures identified on the whole dataset. We tested whether *mastR* can produce highly consistent results across the two different sampling scenarios. We conducted 1,000 random sampling iterations for both strategies to assess the robustness and estimate the error.

#### 2.4.2 Precision, recall and negative control

To evaluate the accuracy, *mastR* was applied to simulated bulk and scRNA-seq data and the identified marker genes were compared with the simulated ground truth. Statistical metrics were computed, namely precision, recall, F1 score and false positives in the negative control of the *mastR* results. The bulk RNA-seq data was simulated using the same method as above. The package *splatter* (Zappia, et al., 2017) was used to simulate scRNA-seq data consisting of 1000 genes with 3000 cells equally divided into 4 groups. For both simulated bulk and scRNA-seq data, groups 1 to 3 each had 2% of all genes being DEGs, while group 4 contained no differentially expressed genes (negative control group). Single cells were aggregated using the *pseudo_samples* function in *mastR* based on group. The recall, precision and F1 score were computed by comparing the *mastR*-identified signatures of groups 1 to 3 with the simulated DEGs. The results for group 4 were used as the negative control to estimate the false positives.

### 2.5 Comparison with published curated signatures

To assess the performance on real-biological data, we used *mastR* to automatically generate a NK cell signature from the DICE dataset (Schmiedel, et al., 2018), and the CRC cell lines data from CCLE (Barretina, et al., 2012) was used to identify the background signal when refining the signature. We compared our *mastR*-derived NK cell signature against 3 published curated NK cell signatures, namely from Crinier et al. (*NK Crinier*) (Crinier, et al., 2018), Cursons et al. (*NK Cursons*) (Cursons, et al., 2019) and Shembrey et al. (*NK Shembrey*) (Shembrey, et al., 2022). Performances were compared by conducting gene-set scoring (using *singscore* (Foroutan, et al., 2018)), gene-set enrichment analysis (GSEA), and precision-recall curve (PRC) on an independent dataset, *pbmc3k.final* from *SeuratData* package (Lab, 2020).

### 2.6 Survival analysis

Survival analysis was conducted on TCGA’s Colon Adenocarcinoma (COAD) dataset with respect to patients’ overall survival (OS) and progression-free interval (PFI), patients were stratified by multiple clinical indicators respectively, including sample type, clinical cancer stage, prior malignancy status, prior treatment status, metastatic (M) stage, gender, expression subtype, microsatellite instability (MSI) status and consensus molecular subtypes (CMS).

Kaplan–Meier survival curves and Cox proportional hazard models were generated using R packages *survival* (Therneau, et al., 2000) and *survminer* (Kassambara, 2021) with standard parameters. A total of 522 samples from 458 patients in TCGA-COAD with valid OS and PFI data were analyzed, and p-value calculated using the log-rank test. These samples were further categorized into 3 groups (low, medium and high), based on the 30^th^ and 90^th^ percentile values of NK score using *singscore* (Foroutan, et al., 2018) with *mastR*-derived NK cell signature.

## 3. Results

### 3.1 *mastR* automatically identifies specific gene expression signatures

We tested *mastR*’s ability to automatically identify a natural killer (NK) cell specific signature from DICE dataset and validate that in an independent immune cell dataset (im_data_6). First, *mastR* summarizes a collection of NK markers from LM7, LM22, and PanglaoDB, along with the collection of gene sets associated with the “NATURAL_KILLER” term in MSigDB into a “pool” of markers. Then using this “pool”, *mastR* automatically generates a set of NK signature genes from 15 immune cell types in the DICE dataset (Schmiedel, et al., 2018). We then compared the expressions of the *mastR*-derived NK signature genes in well-known cell types from both the training and independent datasets, with **Figure 2** showing that the derived NK signature shows consistent performance with high specificity to NK cells. We compared NK cells with stimulated CD8+ T cells and B cells as they are the most similar and dis-similar cells to NK cells respectively. The derived NK cell markers are a subset of the original markers and found to be highly expressed in the NK cells and not well expressed in other cell types (e.g. B cells, CD8+ naïve T cells, **Figure 2A, Supplementary Figure S1**). The derived NK signature can clearly distinguish NK cells from the other cell types as shown by the higher rank scores obtained by scoring each sample in DICE using *singscore* (**Figure 2B**). As can be seen in **Figure 2B**, significantly higher NK scores were demonstrated in NK cells, very clearly differentiating them from other immune cell types (**Supplementary Figure S2)**.

**Figure 2.**
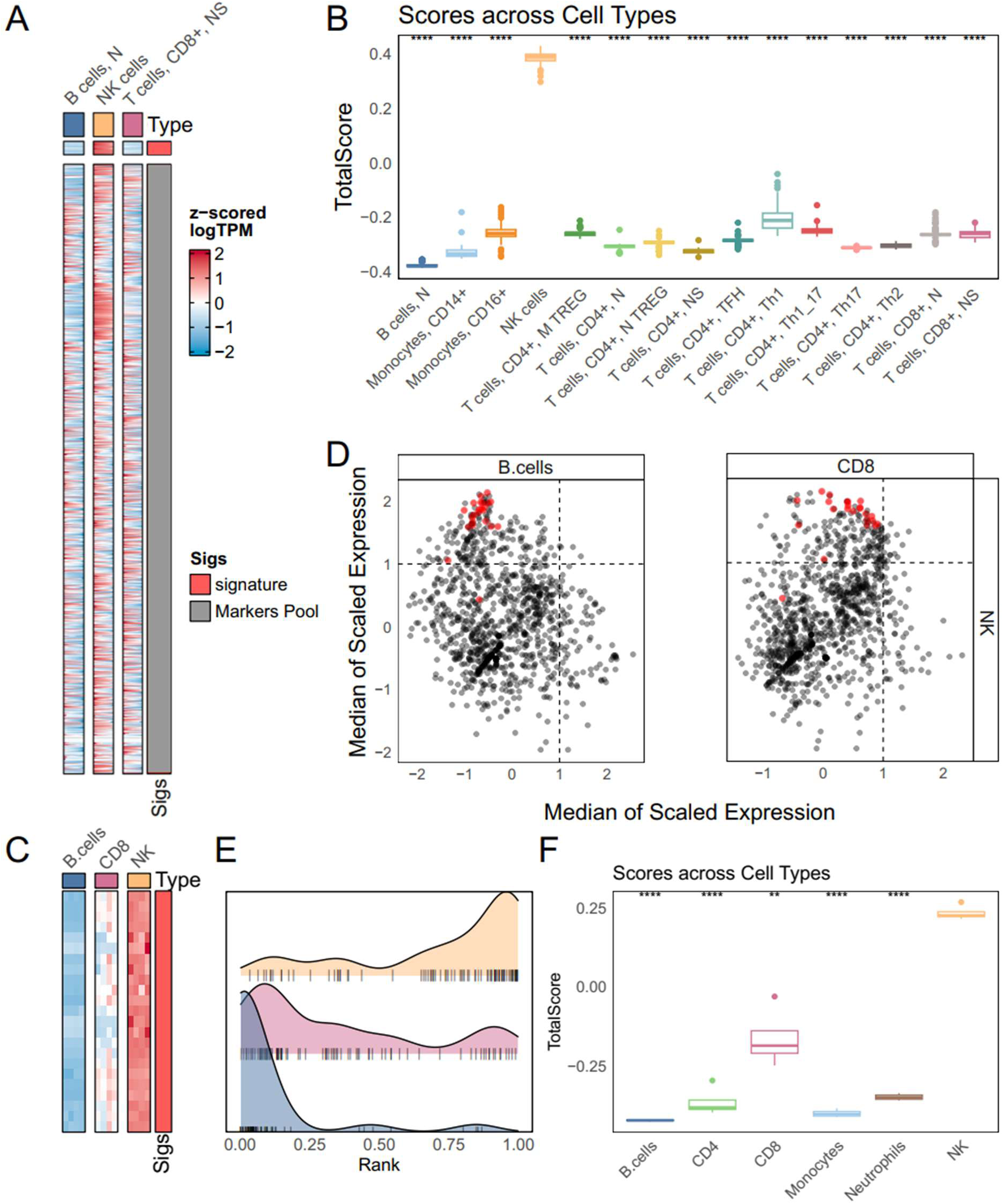
The *mastR*-derived NK cell signature shows good performance on the immune datasets. Performance of *mastR*-derived NK signature on DICE (A-B) and on im_data_6 (C-F). **(A)** Heatmap of scaled log gene expression of all genes in the original markers pool with top section representing the 24 genes from the *mastR*-derived NK signature in DICE; **(B)** Boxplot of ranked scores (using *singscore*) of *mastR*-derived NK signature for the different cell types in DICE; **(C)** Heatmap of scaled log gene expression of *mastR*-derived NK signature in im_data_6; **(D)** Scatter plot of z-scored median gene expression in NK cells (y axis) against z-scored median gene expression in either B cells or CD8+ T cells (x axis) for all genes in markers pool, *mastR*-derived NK signature genes are highlighted in red, with the top-left quadrant represents regions of high NK-specificity; **(E)** Normalized rank density ridges plot of *mastR*-derived NK signature in NK cells (top), B cells (middle) and CD8+ T cells (bottom) respectively; **(F)** Boxplot of ranked scores (using *singscore*) of *mastR*-derived NK signature for the cell types in im_data_6. * M = memory; N = naïve; S = Stimuli in (A) and (B).

We then validated the signature on an independent dataset *im_data_6* (GSE60424 (Linsley, et al., 2014)), which is a sorted bulk RNA-seq data from the blood of healthy individuals (**Figure 2C-F and Supplementary Figure S3**). Our derived NK signature has distinctly higher expression in NK cells compared with CD8 or B cells (**Figure 2C)** consistent with the results on the DICE dataset. This high specificity for NK cells is further illustrated by the normalized median expression biplot of NK cells against other cell types where the signature genes are located in the NK high / (B cells or CD8+ T cells low) quadrant (top-left, red points), distinct from the heterogenous distribution of the original markers in the pool (black points) (**Figure 2D)**. The normalized ranked density barcode of the signature on the NK, CD8 and B cells (**Figure 2E)** suggests an enrichment of our NK signature genes in NK cells but not the other cell types. This enrichment is reinforced via GSEA showing significant over-representation of the signature genes in NK subsets (**Supplementary Figure S3D,** p < 0.01). Finally, the ranked scores of the signature clearly show the effective identification of NK cells from the other cell types (**Figure 2F)**. Taken together, these results suggest that *mastR* can automatically derive a cell type expression signature that is highly specific.

### 3.2 Marker genes identified by *mastR* are accurate and reproducible

We assessed the reliability and robustness of *mastR* approach using a simulation study. Bulk RNA-seq data (100 samples across 4 groups, 1000 genes at 2% DE rate with balanced composition) was simulated as per described in the Methods section. We first identified the signature for each group using *mastR* on the whole simulated dataset, finding all identified signature genes overlap exactly with the ground truth (i.e. simulated differentially expressed genes (DEGs), **Supplementary Figure S4A**. However, in the simulated ground truth data, there is one shared DEG between Group 1 and Group 4 (“Gene463”, colored blue in **Supplementary Figure S4A**) which is not identified and not amongst the *mastR*-derived signatures. This gene has similar high expression in both Group 1 and Group 4 as compared to the rest of the groups (**Supplementary Figure S4B)**. This suggests *mastR* only identifies highly specific genes across the groups, with no false positive markers identified.

We then test the reproducibility of the *mastR* by applying onto sub-sampled datasets with the same group compositions. For each subset, we randomly selected 80% of the samples from each group to generate new group signatures and compared this with the signature derived earlier using the whole dataset using the Jaccard index (JI, Jaccard similarity coefficient). The resulting JI distribution in **Supplementary Figure S4C** (for balanced data) indicates a high similarity with JI > 0.85 (sampling of 1,000), suggesting that *mastR* can stably reproduce the signatures regardless of the size of the dataset. Similarly, we evaluated the sensitivity of *mastR* to varying group sizes using an imbalanced sampling strategy where the sampling probabilities for each group are 0.4, 0.5, 0.7, and 0.8 (groups 1 to 4 respectively). Similar results are obtained as shown in **Supplementary Figure S4C** (for imbalanced data), with similarity JI of > 0.85 (medians of JI, sampling of 1,000). This implies that *mastR* can accurately identify group-specific marker genes regardless of dataset sizes or difference in group composition in a robust manner.

The stability and performance of *mastR* was further evaluated as on the 1000 simulations as described by computing the precision, recall, F1 score and false positives metrics by comparing the derived signatures with the simulated ground truth. Results indicate that *mastR* achieves excellent performance with a F1 score of 0.97 ± 0.04, 100% precision, a recall of 0.95 ± 0.06 and low false positives rate of 0.01 ± 0.07 when comparing *mastR*-derived signatures with all simulated DEGs (**Table 3**, “all DEGs”). A smaller recall was noted compared to the F1 score, suggesting some “DEGs” that may not be identified by *mastR*. These specific “DEGs” were further assessed and found to be highly specific for more than just one group, as was the case for “Gene463” (**Supplementary Figure S4B)**. Thus, we refined the comparison to only compare the derived signatures with only the simulated DEGs that’s unique to each group. In this case, the performance improved as expected with an F1 score of 0.99 ± 0.03 and a recall of 0.99 ± 0.05 (**Table 3**). The results demonstrate *mastR* can accurately identify marker genes with high specificity, particularly for those unique marker genes to each group in the data.

**Table 3.** Performance of mastR on simulated bulk RNA-seq data.

| metric | mean | sd | DEGs |
| --- | --- | --- | --- |
| F1 Score | 0.97 | (0.04) | All |
|  | 0.99 | (0.03) | Unique |
| Precision | 1.00 | (0) | All |
|  | 1.00 | (0) | Unique |
| Recall | 0.95 | (0.06) | All |
|  | 0.99 | (0.05) | Unique |
| False Positives | 0.01 | (0.07) |  |
| Running Time | 2.91 | (0.39) |  |

Although *mastR* was originally designed for bulk RNA-seq data, we further assessed its applicability on scRNA-seq data simulated using the *splatter* (Zappia, et al., 2017) package as described in the **Methods** section (3000 cells across 4 groups, each 1000 genes at 2% DE rate with balanced composition). The single cells were aggregated (pseudo-bulked) and performance evaluated against commonly used scRNA-seq package *Seurat* (Hao, et al., 2021) for 1,000 simulations. Results as shown in **Table 4**, suggests that *mastR* achieves better performance than *Seurat* on simulated scRNA-seq across all the performance metrics, with higher accuracy and lower false positives. Importantly, it should be noted that *mastR* is significantly more computational efficient (∼ 1.29 seconds on average) compared with Seurat (∼ 60 seconds on average) which is important for real life applications. Therefore, *mastR* can handle large-scale single cell datasets with complex scenarios at a lower computational cost, showing a significant speed advantage and high accuracy.

**Table 4.**
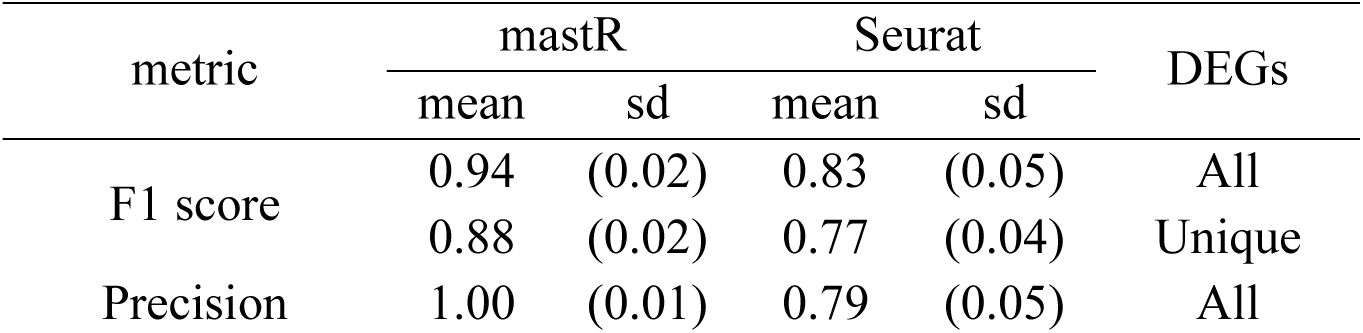
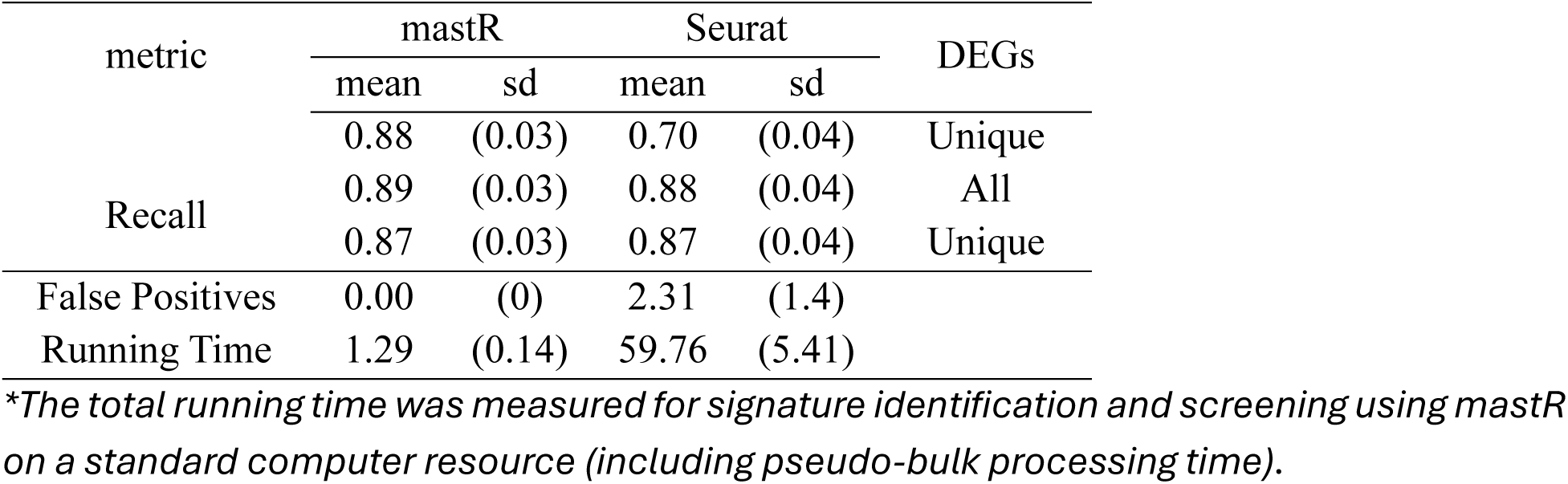
Performance of mastR and Seurat on simulated scRNA-seq data.

### 3.3 The *mastR*-derived gene expression signature performs as well if not better than manually curated signatures

We further explored the performance of our automatically generated NK signature (*NK mastR*) compared to the published manually curated signatures, using an independent dataset *pbmc3k.final* (Lab, 2020). Three published NK cell signatures (Crinier, et al., 2018; Cursons, et al., 2019; Shembrey, et al., 2022) were collated for this study, namely from Crinier *et al (NK Crinier*), Cursons *et al* (*NK Cursons*) and Shembrey *et al* (*NK Shembrey*). All 4 signatures are able to distinguish NK cells from all other cell types when comparing normalized rank scores, with CD8+ T cells most closely resembling NK cells (**Figure 3A**). However, for *NK Crinier* and *NK Cursons*, the majority of CD8+ T cells exhibit a higher score than the lowest score of the NK cells (arrows in **Figure 3A**), indicating a higher degree of similarity between these 2 signatures. This is also shown in the mean differences of the ranked scores between NK cells and CD8+ T cells, where both *NK Crinier* and *NK Cursons* have the smallest differences (< 0.40). This observation is further validated using the area under the precision-recall curve (AUPRC) in **Figure 3B**, where only *NK mastR* and *NK Shembrey* achieved an AUPRC of > 0.90. This suggests that the automatically generated NK signature is performing as well if not better than manually curated signatures in distinguishing between cell types.

**Figure 3.**
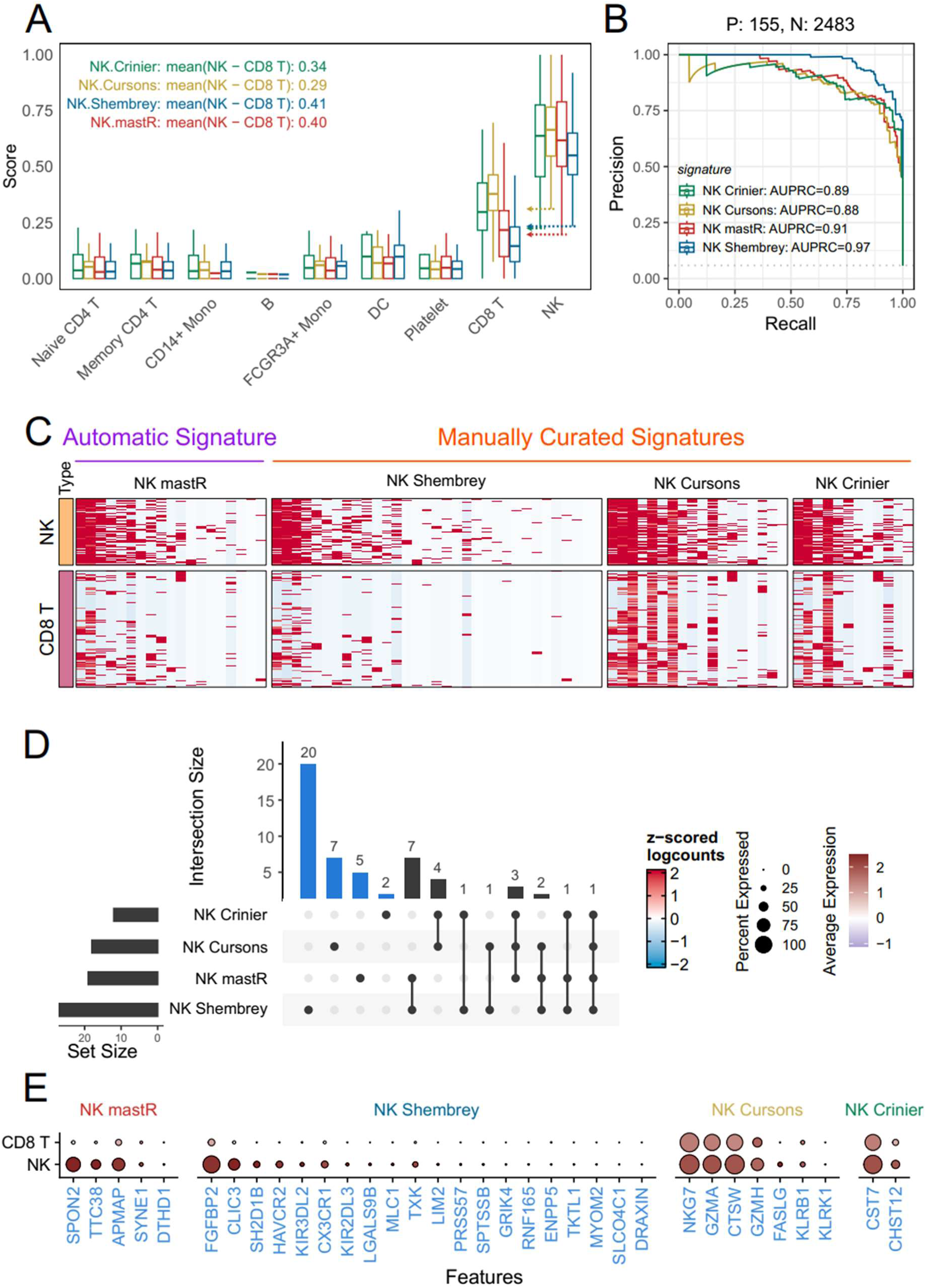
The *mastR*-derived NK signature performance as well as published curated signatures. Comparisons of signatures performance using independent scRNA-seq data (pbmc3k.final). **(A)** Boxplot of scaled ranked scores (using *singscore*) on the 4 NK signatures of interest with the mean differences in ranked scores between NK and CD8 T cells calculated and shown (inset). The minimum scores of NK cells for each signature are shown as a colored arrow for each boxplot; **(B)** Precision-recall curve (PRC) of the NK signatures with the individual area under curve (AUC) computed; **(C)** Heatmap of scaled log gene expression of the 4 NK signatures for both CD8+ T cells and NK cells; **(D)** Overlap between the 4 NK signatures. Blue bars indicate numbers of unique genes for each signature; **(E)** Dot plot of the average gene expression of the unique signature genes in each signature (blue bars in *D*) for CD8+ T cells and NK cells, color represents average expression and size represents expressed percentage. * Note that only genes in pbmc3k.final dataset are shown and used in subsequent analysis.

We then access the gene signatures at the individual gene level where it is noted that most of the signature genes have high expression in NK cells while less so for CD8 T cells (**Figure 3C)**. However, it should be noted that both *NK Cursons* and *NK Crinier* also have relatively higher expression in CD8 T cells compared to the other signatures. On the other hand, a significant number of *NK Shembrey* signature genes also have low expression in both NK and CD8 T cells. By comparing the overlapping genes between the signatures in **Figure 3D**, it is clear that each signature has a set of unique genes (in blue) with only *KLRF1* (an essential gene expressed in most NK cells) common amongst all 4 signatures. We assessed the performance of these unique genes by comparing their average expression within NK cells or CD8+ T cells (**Figure 3E)**. Most of the unique genes in our NK signature is highly specific to NK cells while exhibiting low expression and proportions in CD8+ T cells, while other signatures are either specific to NK and CD8+ T cells (*NK Cursons* and *NK Crinier*) or have many unique genes non-specific to NK cells (*NK Shembrey*). These results are also found to hold up when comparing across all cell types (**Supplementary** Figure 5), confirming the ability of *mastR* to identify novel and highly specific markers.

### 3.4 The *mastR*-derived signature as a potential indicator of clinical outcomes

We further assess how our derived NK signature can be applied to clinical applications. Here we apply our NK signature to the TCGA-COAD dataset (**Supplementary Figure S6-S8**) to assess how well it can delineate clinical outcomes. In this analysis, the colorectal-associated cell lines data from CCLE (Barretina, et al., 2012) was used as the reference data to remove colon specific signature genes (i.e. highly expressed with SNR < 1 in CRC-related cell lines). Each sample was then rank scored and the subsequent NK scores were compared across different clinical indicators. The results suggest that NK signature has no significant correlation with prior malignancy status or gender (**Supplementary Figure S6**). However, significantly different NK signature ranked scores were observed across various clinical indicators, indicating different extent of NK cell infiltration under different conditions. In particular, amongst the consensus molecular subtypes (CMS), CMS 1 subtype has a significantly higher NK score than the other subtypes (**Figure 4A**). This is consistent with the findings from the study by Guinney and colleagues (Guinney, *et al*., 2015) where it was suggested that CMS 1 subtype tumors have a strong immune response and higher NK cell activity. Our results also indicate a significantly elevated NK ranked score in microsatellite instability (MSI) high samples compared to others (**Figure 4B**). This potentially suggests increased NK cell infiltrations in MSI high tumors which agrees with a previous finding by Lanuza *et al*. (Lanuza, et al., 2022).

**Figure 4.**
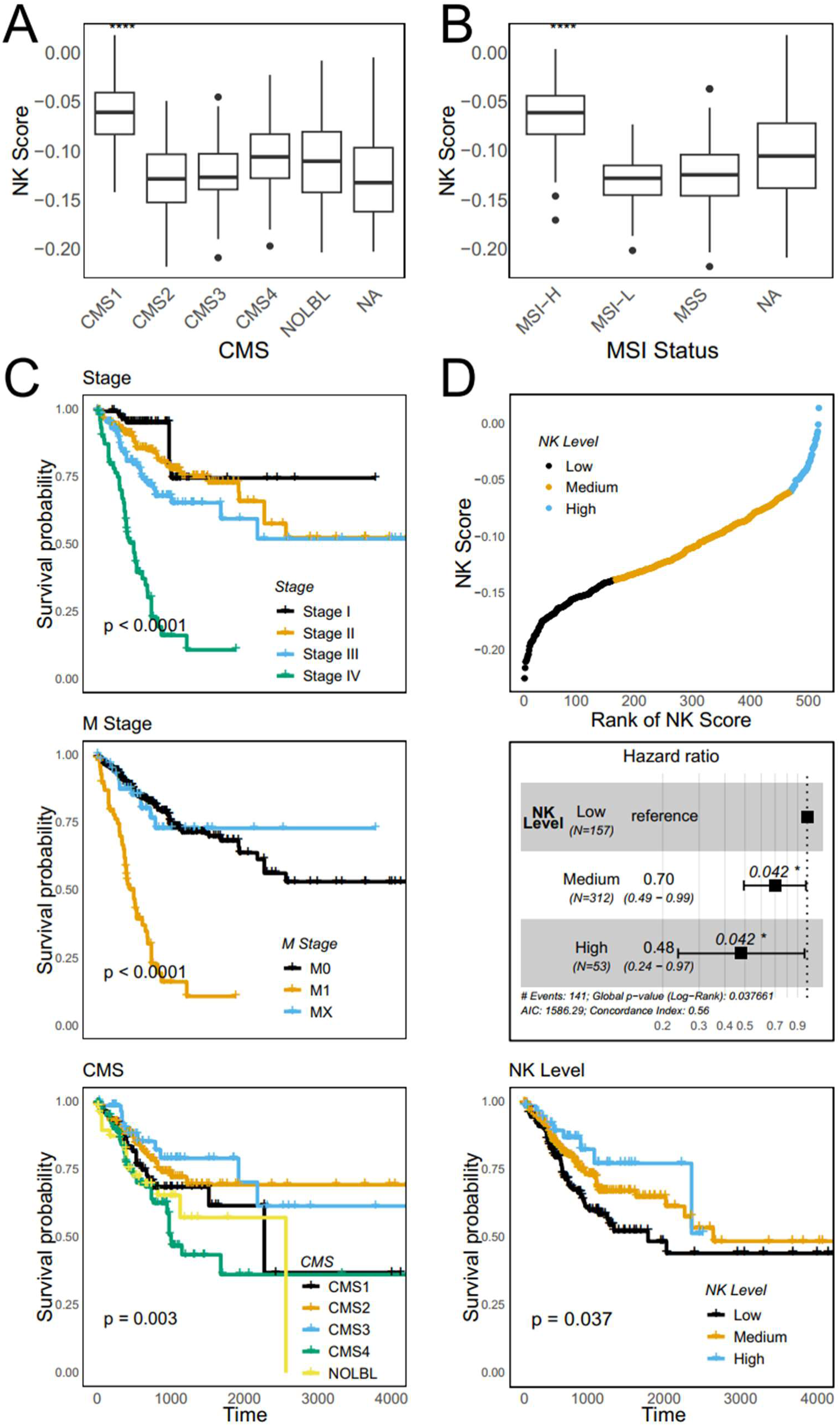
Application of NK signature on TCGA-COAD dataset. **(A)** Boxplot of NK ranked scores (using *singscore*) across (A) consensus molecular subtypes (CMS) and **(B)** microsatellite instability (MSI) groups, with significance calculated by t test; **(C)** Survival analysis of progression-free interval (PFI) across clinical stages (top), metastatic (M) stages (middle) and CMS groups (bottom) with log-rank test p value; **(D)** Samples are categorized into 3 groups based on the 30^th^ and 90^th^ percentile of NK ranked scores with samples ordered by NK ranked scores (top); forest plot of Cox proportional hazards model of PFI on NK score groups (middle); and survival analysis of PFI on NK score groups with log-rank test p value (bottom). Survival analyses are conducted using packages *survival* and *survminer* with default parameters on 522 samples from 458 patients.

Taking the clinical indicators showing significant differences in NK scores, survival analyses were performed where we found that overall survival (OS) is significantly associated with clinical disease stage, M (metastatic) stage and prior treatment status (**Supplementary Figure S7**), while progression-free interval (PFI) is significantly associated with clinical stage, M stage and CMS groupings (**Supplementary Figure S8**). These results suggest that patients at a more advanced disease stage or M stage (**Figure 4C**, top & middle), show poorer outcomes in terms of both OS and PFI, and both conditions are associated with a low NK score. Interestingly, CMS survival analysis presents a different trend for PFI, where CMS1 (which had the highest NK ranked scores amongst the CMS subtypes) (**Figure 4A**) shows a poor PFI similar with that of CMS4 (**Figure 4C**, bottom). This was also observed in the study by Guinney *et al* (Guinney, et al., 2015) which also concluded that both CMS1 and CMS4 presents the worst relapse-free outcomes. This suggests that CMS1 is a heterogenous cancer subtype where survival outcomes are associated with multiple factors and not solely due to NK cell infiltration. Thus, to investigate the contribution of NK signature ranked score to CRC survival outcomes, the samples were categorized into 3 groups based on the 30^th^ and 90^th^ percentile of NK ranked scores: depicted as low, medium and high respectively (**Figure 4D**, top). Subsequent survival analysis for PFI (**Figure 4D**, middle & bottom) shows a significant association between NK ranked scores and the patients’ PFI, where a higher NK score is associated with lower risk. Taken together, these results suggest that *mastR*-derived NK signature can potentially be applied to clinical data as a indicator of clinical outcomes.

## 4. Discussion

Here we introduce our tool, *mastR*, an innovative R/Bioconductor package to automate biological marker identification. Leveraging the statistical robustness of *edgeR* and *limma*, and taking rank product-based scores, *mastR* automates the identification of group-specific markers from bulk RNA sequencing data. A key feature distinguishing *mastR* from standard marker selection methods is the reduction of background microenvironment signal using appropriate reference data. This capability enhances the application of *mastR* in scenarios where the effects of background signals confound the marker signal, and in cases which require careful consideration, such as in contexts involving disease-specific, tissue-specific or organ-specific datasets. Though in certain scenarios, it is necessary to obtain markers from multiple datasets to ensure robustness and confidence. *MastR* extracts the markers for each dataset separately and aggregates the results from all datasets, instead of employing a meta-analysis based on data integration. The advantage of this approach is that it avoids any over-normalization or mis-correction that may occur when integrating datasets with strong batch variances together into a common dataset, leading to a robust and conserved signature for the datasets (Rau, et al., 2014). However, in some cases data integration meta-analysis might be more suitable, depending on the data and researcher preference and particularly when the dataset lacks replicates.

In this study, we evaluated *mastR’s* performance on both simulated and biological datasets, as well as compared of the performance of the *mastR*-derived NK cells signature against published NK signatures, with *mastR* demonstrating high accuracy and robustness. While the focus of this study was to evaluate *mastR* for marker identification in bulk RNA-seq data, it is important to acknowledge the potential application of *mastR* on scRNA-seq datasets. We compared *mastR’s* performance with *Seurat*, one of the main statistical packages designed for scRNA-seq data. Impressively, *mastR* performed better than *Seurat* and had significantly reduced computation time.

While not explicitly discussed, in some marker identification studies, the presence of two or more closely related groups in the data may pose challenges for the identified markers to be effective in distinguishing these groups (Burel, et al., 2022). In such cases, *mastR* allows the preservation of the top features from specific DE comparisons for further refinement and performing rank product scoring by utilizing the underlying statistics (e.g., logFC, adjusted p values). This increases the discriminative power of signature between similar groups and improve signature identification based on statistics.

Although this paper focuses primarily on transcript-based data, *mastR* can theoretically be applied to all types of multi-omics data. We aim to explore this in the future and to validate the performance of *mastR* using experimental data across diverse omics types to improve application and generalizability across a range of research contexts. However, several challenges remain despite the best efforts of *mastR*. Firstly, *mastR* does not work when there is no suitable sample variable to aggregate single cells (e.g., fewer than 2 replicate pseudo-samples within a group), in which cases specific methods for those scRNA-seq datasets are still needed. Secondly, by default *mastR* utilizes intersection of DEGs across DE comparisons. This approach, while effective, may result in cases where there are no or limited numbers of intersecting markers, particularly when dealing with many groups. In this situation, it might be necessary to either merge some of the groups (less groups) or select the “union” assemble method in *mastR* instead of the default.

## Acknowledgements

We would like to thank Ashley Weir for reviewing and providing constructive feedback for this manuscript.

## Data and code availability

All datasets underlying this article are available from public data repository or publications as described in **Table 1**. The code used for the performing the described analyses is available from the corresponding author upon reasonable request. The *mastR* package is publicly available on Bioconductor (version ≥ 3.17) at https://bioconductor.org/packages/release/bioc/html/mastR.html.

