## Supplementary Figures for "mastR: Marker Automated Screening Tool for multi-omics data"

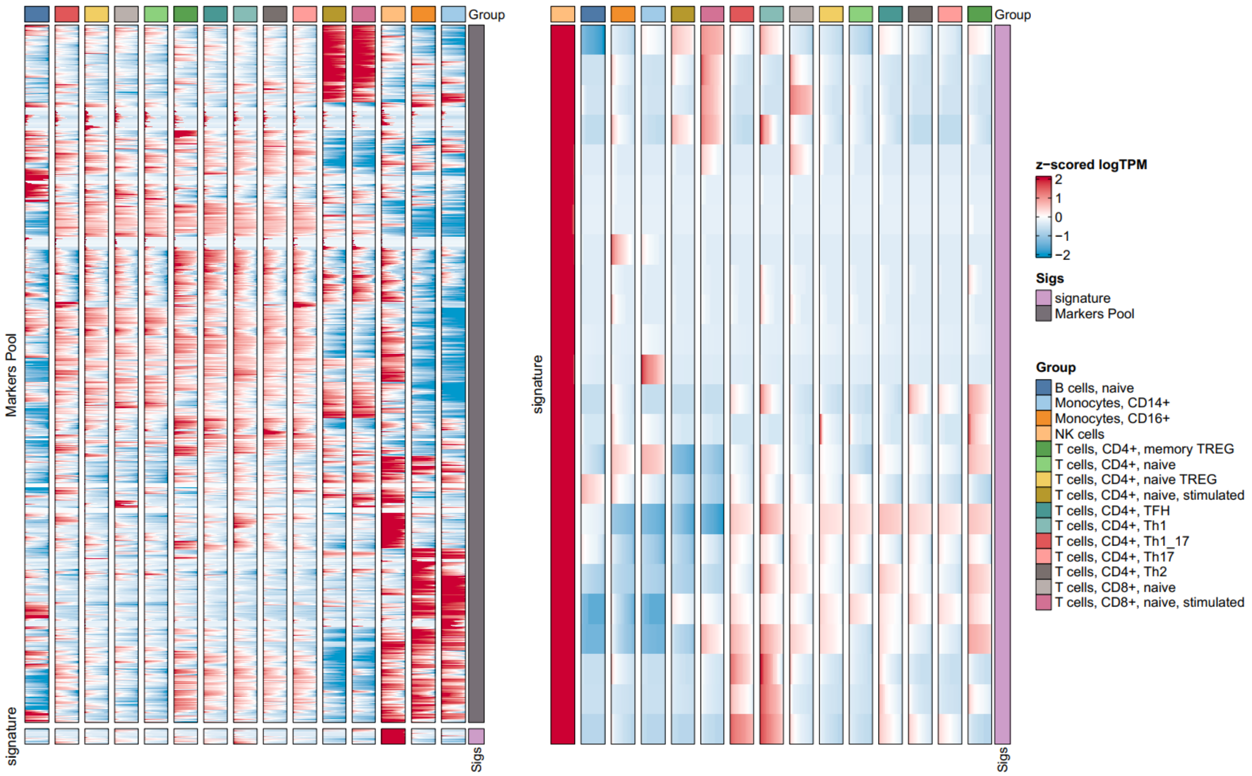


**Supplementary Figure 1. *mastR*-derived NK signature is enriched in NK cells.** Heatmap of scaled gene expression (logTPM) of NK cell marker genes in DICE. (Left) Heatmap of the whole original set of markers across all cell types in DICE. (Right) Heatmap of the *mastR*-derived NK signature genes across all cell types in DICE.

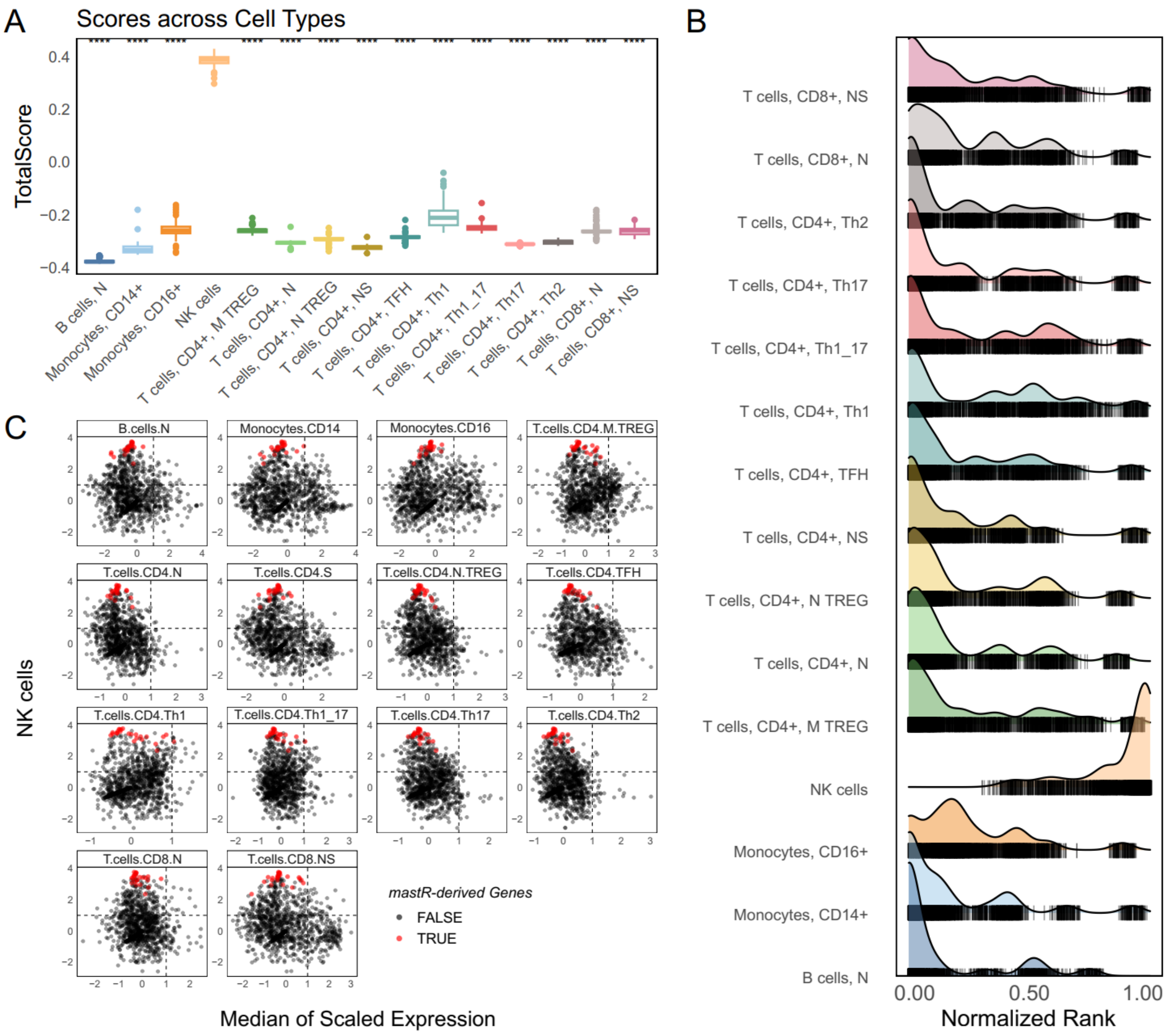


**Supplementary Figure 2. The *mastR*-derived NK signature is highly specific to NK cells in DICE.** Overall performance of NK signature in DICE dataset. **(A)** Boxplot of ranked scores (using *singscore*) based on derived NK signature across cell types in DICE; **(B)** Normalized rank density ridges plot of derived NK signature for each cell type; **(C)** Scatter plot of z-scored median gene expression in NK cells (y axis) against z-scored median gene expression in other cell types (x axis) for all the genes in the original set of markers. The *mastR*-derived NK signature genes are highlighted in red, top-left quadrant represents region of high NK-specificity. * M = memory; N = naïve; S = Stimuli in (A), (B) and (C).

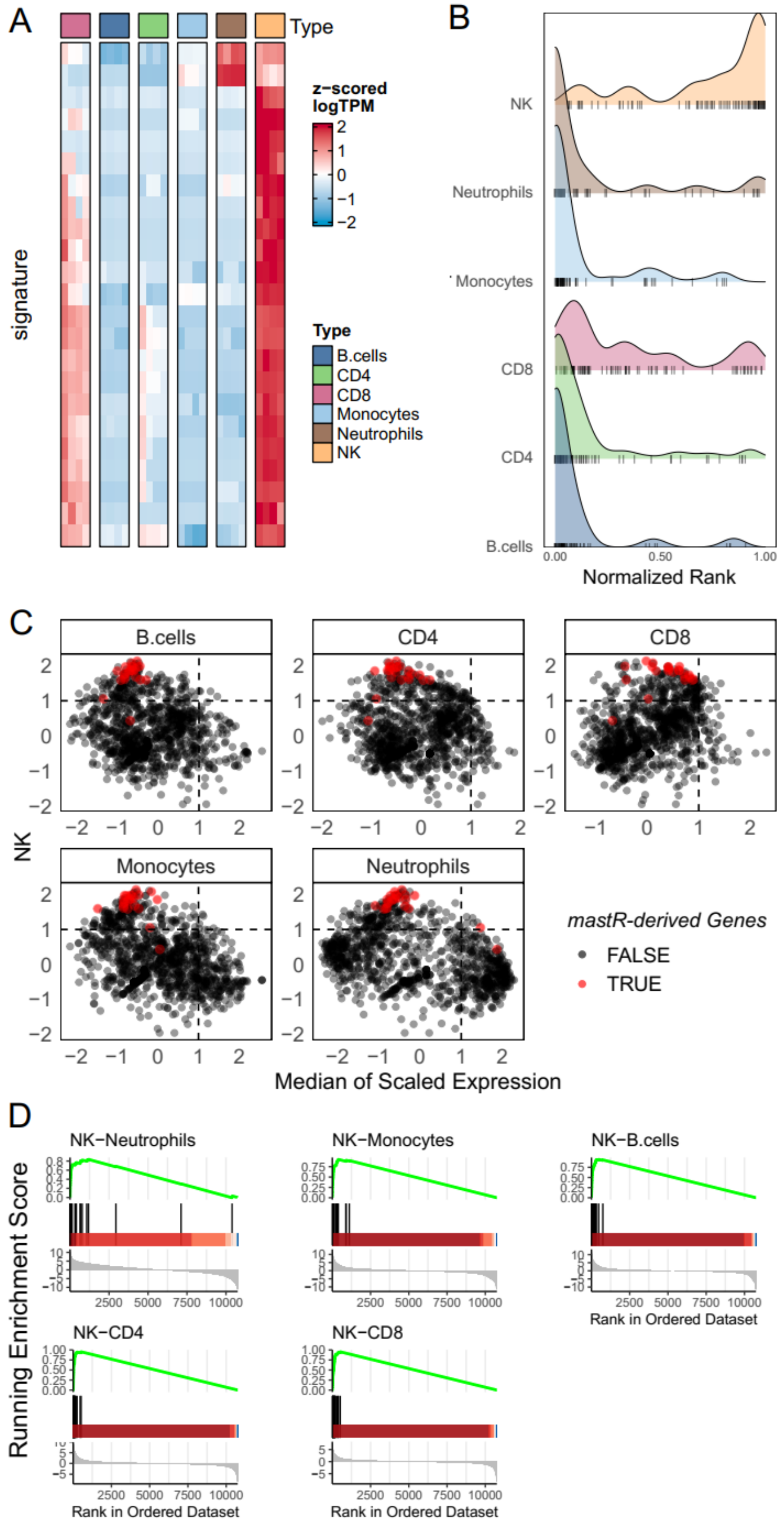
A
Type
signature
z-scored
logTPM
2
1
0
−1
−2
Type
B.cells
CD4
CD8
Monocytes
Neutrophils
NK
B
NK
Neutrophils
Monocytes
CD8
CD4
B.cells
0.00
0.50
1.00
Normalized Rank
C
B.cells
CD4
CD8
Monocytes
Neutrophils
NK
Median of Scaled Expression
mastR-derived Genes
FALSE
TRUE
D
NK−Neutrophils
NK−Monocytes
NK−B.cells
NK−CD4
NK−CD8
Running Enrichment Score
Rank in Ordered Dataset
2500
5000
7500
10000

**Supplementary Figure 3. The *mastR*-derived NK signature performs well when applied on the independent dataset *im_data_6*.** Performance overview of derived NK signature in *im_data_6*. **(A)** Heatmap of scaled log gene expression of NK signature across cell types in *im_data_6*; **(B)** Normalized rank density ridges plot of NK signature for each cell type in *im_data_6*; **(C)** Scatter plot of z-scored median genes expression in NK cells (y axis) versus other cell types (x axis) in *im_data_6*, *mastR*-derived NK signature genes are highlighted by red, left-top quadrant represents high NK-specificity; **(D)** Gene-set Enrichment Analysis (GSEA) results of *mastR*-derived NK signatures for the comparisons of NK vs other cell types in im_data_6 presented as running enrichment scores against barcode plots (all $p < 0.01$).

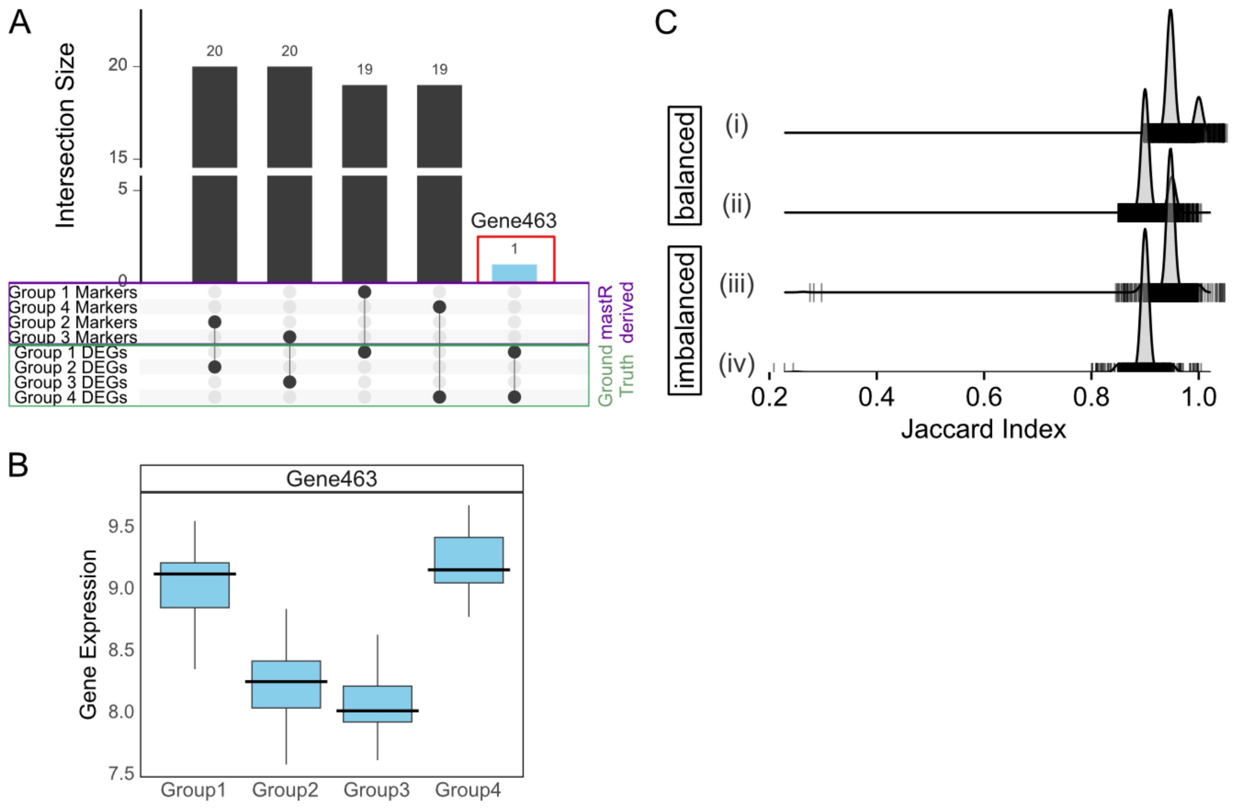


**Supplementary Figure 4. The *mastR* is able to accurately and robustly identify DE markers. (A)** Upset plot of *mastR*-derived group signatures ("Group 1-4 Markers") and simulated up-regulated DEGs ("Group 1-4 DEGs", the ground truth). Note *mastR* did not identify gene "Gene463" which is commonly DE in both Group 1 and 4 of the DEGs ground truth groups (in blue); **(B)** Boxplot of the expression (logCPM) for the simulated up-regulated DEG "Gene463" shared between Group 1 and 4; **(C)** Ridges plot of Jaccard index (JI) across 1,000 random samples of the simulated bulk RNA-seq data. Ridges plot of JI for balanced (top, 80% of each group sampled to obtain new signature) and im-balanced (bottom, 40%, 50%, 70%, and 80% of groups 1 to 4 sampled respectively) sampling strategy on simulated bulk RNA-seq data. The curves (i) & (iii) represent JI between the *mastR*-derived new signature (from the sub-samples) and the derived original signature (from the whole dataset), and (ii) & (iv) represent JI between the *mastR*-derived new signature (from sub-samples) and the simulated DEGs (ground truth) (ii & iv).

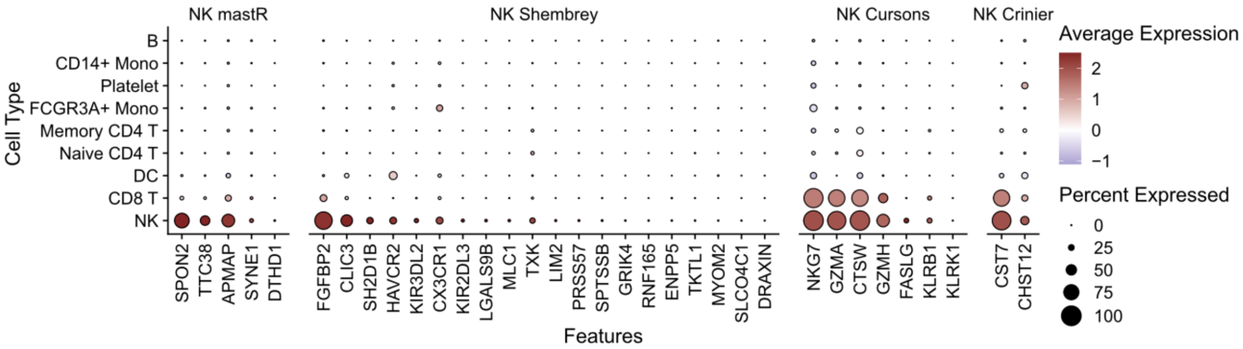


**Supplementary Figure 5. Dot plot of average expression of the unique genes for each NK signature across the cell types in pbmc.3k.final**. Color represents average expression, size represents percent of expressed cells.

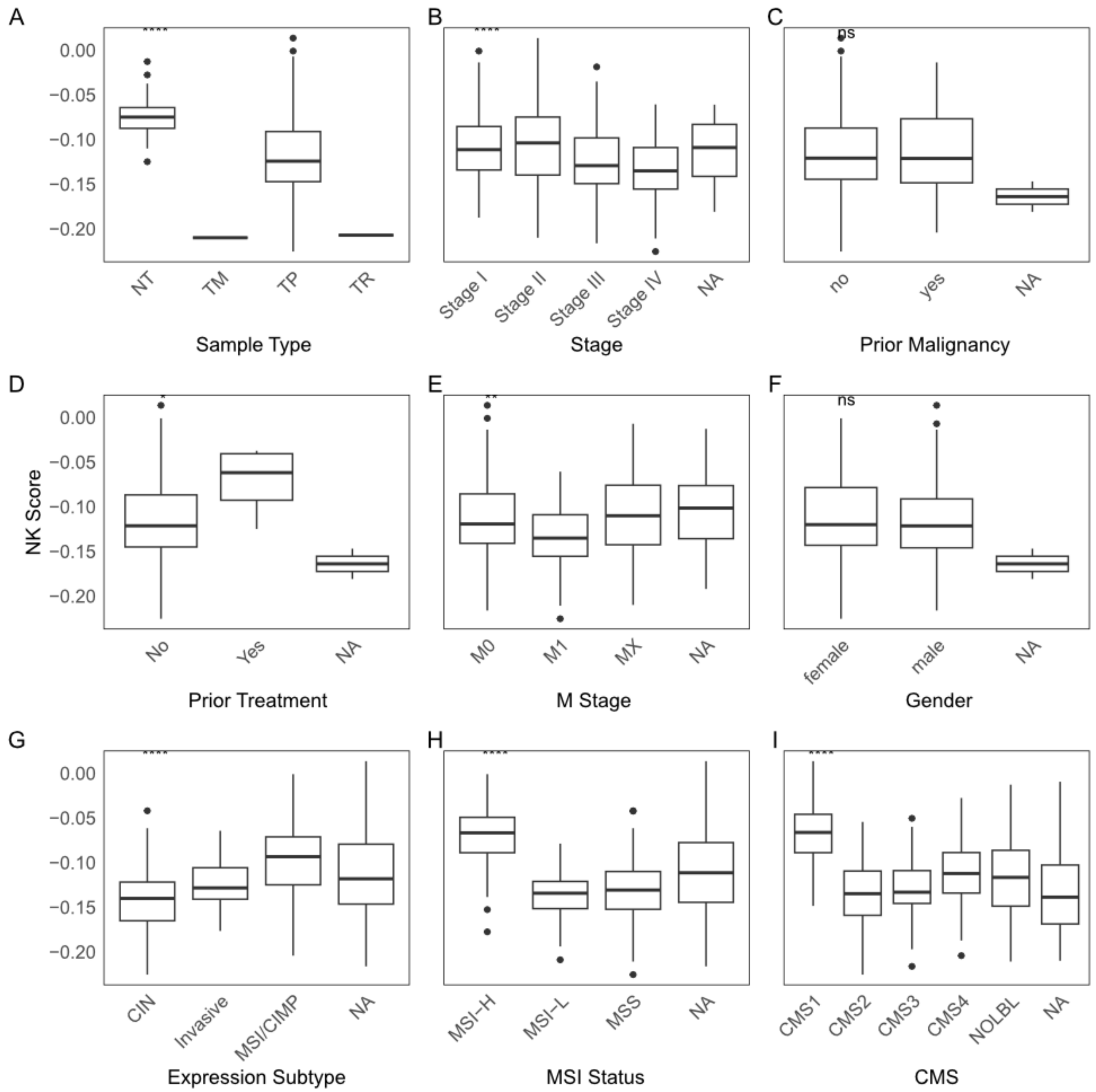


**Supplementary Figure 6. NK signature score shows significant differences between clinical indicators.** Boxplot of the ranked scores of derived NK signature for different clinical conditions in TCGA-COAD dataset. NK ranked scores was plots based on (A) tissue types, (B) cancer stages, (C) malignancy status, (D) treatment status, (E) metastatic (M) stages, (F) gender, (G) expression subtypes, (H) microsatellite instability (MSI) status and (I) consensus molecular subtypes (CMS). * p-value ≤ 0.05; ** p-value ≤ 0.01, *** p-value ≤ 0.001, **** p-value ≤ 0.0001, ns p-value > 0.05.

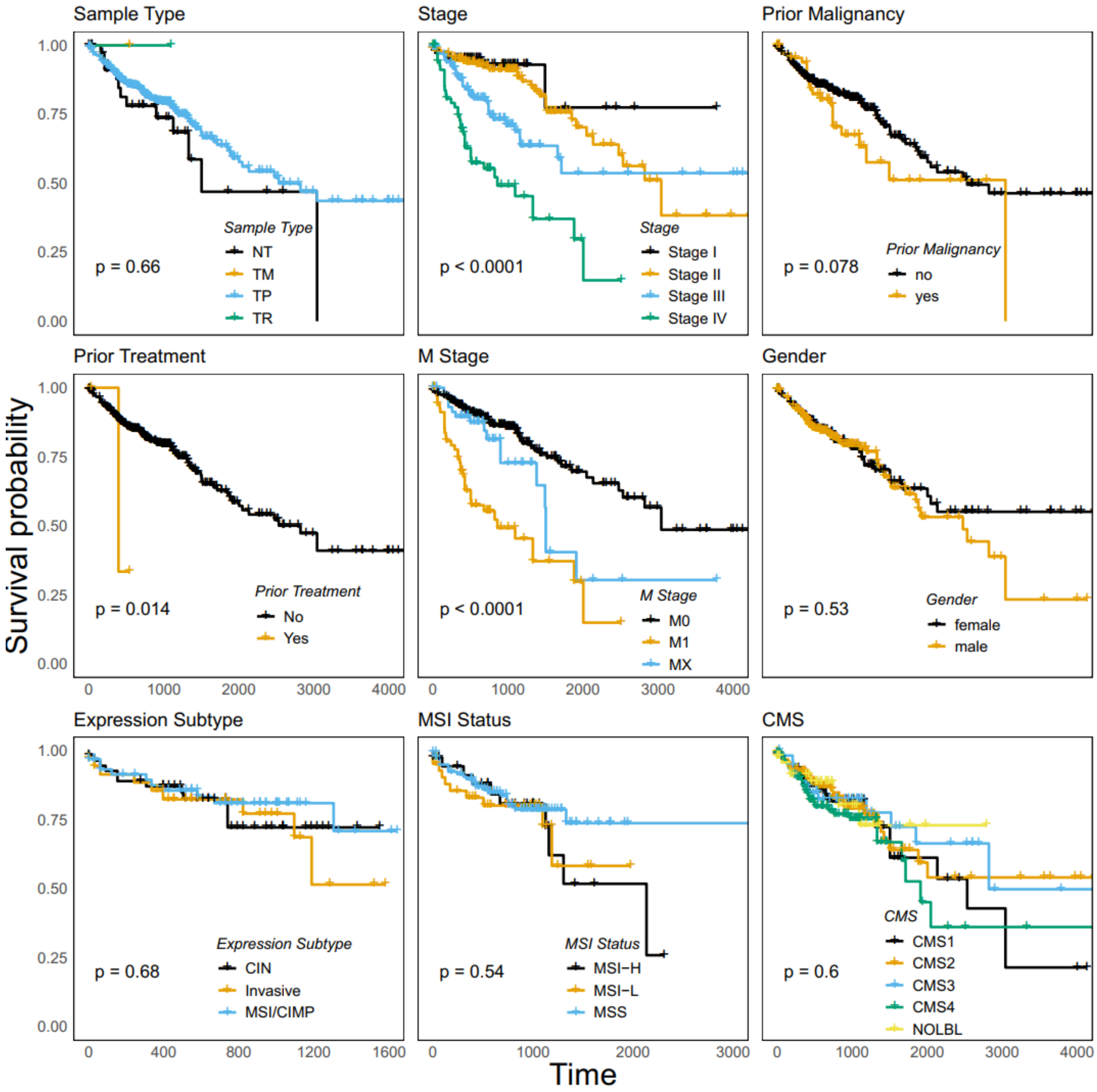


**Supplementary Figure 7. Survival analysis for overall survival (OS) across clinical indicators in TCGA-COAD dataset.** Survival analysis was conducted using packages *survival* and *survminer* using default parameters, a total of 522 samples from 458 patients were analyzed, with log-rank test p value shown.

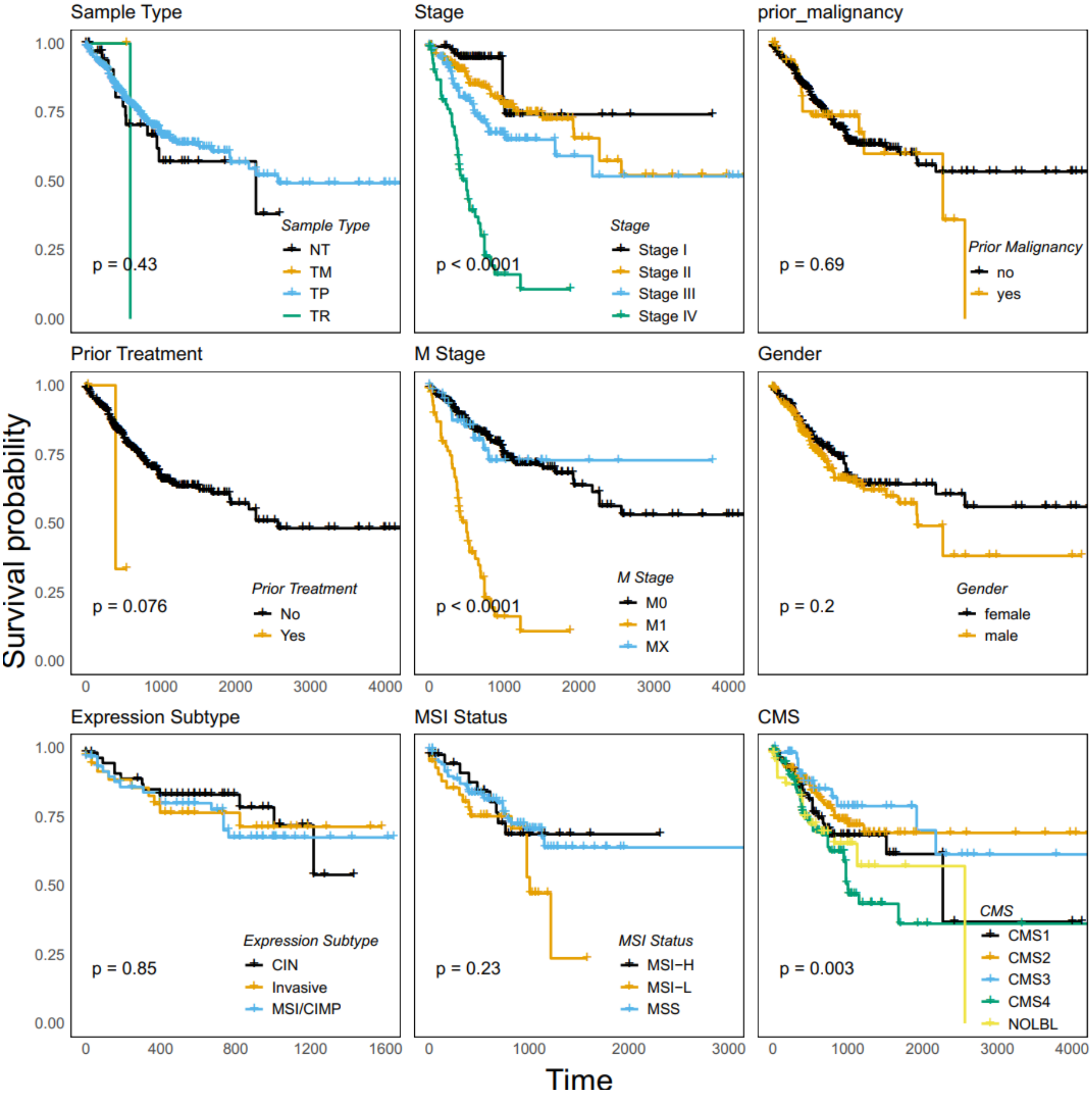


**Supplementary Figure 8. Survival analysis for progression-free interval (PFI) across clinical indictors in TCGA-COAD dataset.** Survival analysis was conducted using packages *survival* and *survminer* using default parameters, a total of 522 samples from 458 patients were analyzed, with log-rank test p value shown.
